# An Automated Approach to the Quantitation of Vocalizations and Vocal Learning in the Songbird

**DOI:** 10.1101/166124

**Authors:** David G. Mets, Michael S. Brainard

**Affiliations:** Department of Physiology, University of California, San Francisco, CA 94158; Center for Integrative Neuroscience, University of California, San Francisco, CA 94158; Howard Hughes Medical Institute, University of California, San Francisco, CA 94158.

## Abstract

Studies of learning mechanisms critically depend on the ability to accurately assess learning outcomes. This assessment can be impeded by the often complex, multidimensional nature of behavior. We present a novel, automated approach to evaluating imitative learning that is founded in information theory. Conceptually, our approach estimates the amount of information present in a reference behavior that is absent from the learned behavior. We validate our approach through examination of songbird vocalizations, complex learned behaviors the study of which has provided many insights into sensory-motor learning in general and vocal learning in particular. Historically, learning has been holistically assessed by human inspection or through comparison of specific song features selected by experimenters (e.g. fundamental frequency, spectral entropy). In contrast, our approach relies on statistical models that broadly capture the structure of each song, and then uses these models to estimate the amount of information in the reference song but absent from the learned song. We show that our information theoretic measure of song learning (contrast entropy) is well correlated with human evaluation of song learning. We then expand the analysis beyond song learning and show that contrast entropy also detects the typical song deterioration that occurs following deafening. More broadly, this approach potentially provides a framework for assessing learning across a broad range of similarly structured behaviors.

**Author Summary:** Measuring learning outcomes is a critical objective of research into the neural, molecular, and behavioral mechanisms that support learning. Demonstration that a given manipulation results in better or worse learning outcomes requires an accurate and consistent measurement of learning quality. However, many behaviors (e.g. speech, walking, and reading) are complex and multidimensional, confounding the assessment of learning. One behavior subject to such confounds, vocal learning in Estrildid finches, has emerged as a vital model for sensory motor learning broadly and human speech learning in particular. Here, we demonstrate a new approach, founded in information theory, to the assessment of learning for complex high dimensional behaviors. Conceptually, we determine the amount of information (across many dimensions) present in a reference behavior and then assess how much of that information is present in the resultant learned behavior. We show that this measure provides an accurate, holistic, and automated assessment of vocal learning in Estrildid finches. Potentially, this same approach could be deployed to assess shared content in any multidimensional data, behavioral or otherwise.

## Introduction

Songbird vocal learning shares many parallels with speech learning[1] and is a powerful, tractable model system for elucidating neural and behavioral mechanisms underlying vocal control and vocal learning[2]. Birds, like humans, learn vocalizations early in life through exposure to the vocalizations of an adult ‘tutor’ followed by a period of practice that eventually results in typical adult vocalizations that require auditory feedback for maintenance[1]. Song is composed of discrete units of sound (syllables) organized into higher order sequences[3]. In the finch species examined here, a given bird’s song comprises about 5-10 categorically distinct syllable types, with these distinct types defined by their unique spectro-temporal structure (Fig. 1A). Hence, an individual bird’s song can be described as a specific set of categorically distinct syllable types (that can be labeled ‘A’, ‘B’, ‘C’ and so on).

**Fig 1.**
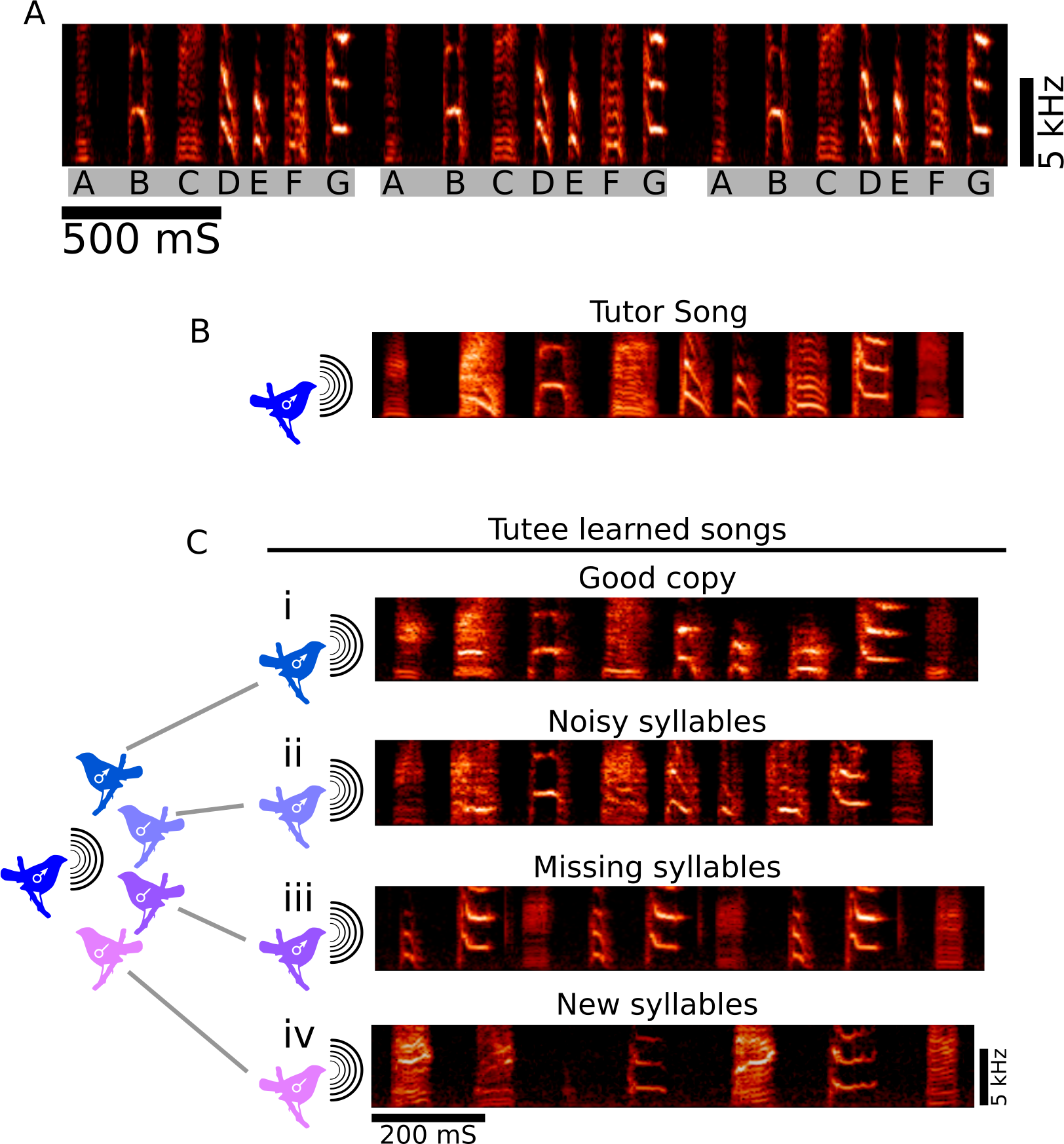
Quantification of song learning is complicated by variety in both learning and failure to learn. (A) Typical sample of song from an adult Bengalese finch. Song is composed of a set of categorically distinct syllable types (labeled ‘A’, ‘B’, ‘C’…) that are organized into larger, repeated, sequences (gray bars). Both the spectral structure of syllables and their sequencing are learned features of song. Hence, song is a complex, high dimensional behavior that differs across individuals. (B) Song of an adult male ‘tutor’ and (C) songs of four juvenile ‘tutees’ that were all exposed to the same tutor song, illustrating variation in the quality of song learning. (Ci) Song from a tutee that learned the spectral content of the tutor song well, producing a song with accurate copies of all syllables. (Cii) Song from a tutee that copied all syllables, but with noisier versions than those present in the tutor song. (Ciii) song from a tutee that failed to copy some of the syllables from the tutor song. (Civ) song from a tutee that included ‘new syllables’ that were not clearly present in the tutor song.

Qualitatively, learning (and failure to learn) can occur in different ways (Fig. 1B-C)[4]. For example, juvenile ‘tutees’ could learn to produce all distinct syllable types present in an adult ‘tutor’ song, but the spectral content of the syllables might be imperfect or noisy (Fig. 1Ci-Cii), while other tutees might completely fail to learn some syllables (Fig. 1Ciii), and still others might improvise new syllables (Fig. 1Civ).

Because of these complexities, many studies have relied on human evaluation of song similarity and learning [5–8]. Indeed, human scorers can provide a useful ‘holistic’ assessment of similarity between complex behaviors, such as songs, which integrates across many stimulus dimensions. However, human scoring suffers from several problems including 1) it requires scorers to be trained on species-specific vocalizations, and analysis of different vocalization types often requires new training, 2) correspondingly, scores are potentially inconsistent over time and across different evaluators, and 3) human scoring is labor intensive and does not readily scale to the size of relevant datasets, which, in the case of bird song, can include many individuals and thousands of vocalizations per day.

More recent attempts at quantification of song similarity have focused on assessing learning based on specific, reliably and automatically measured features of song (e.g. the fundamental frequency of a specific syllable, or song entropy)[9–12]. In these approaches, selected samples of songs are decomposed into sets of feature values[13] and song similarity is then evaluated as the similarity between weighted feature sets associated with those samples. These approaches can overcome some of the inter-evaluator variability associated with human scoring, and additionally enable useful assessment of how specifically analyzed features such as the ‘pitch’ or ‘noisiness’ of syllables differ across songs and conditions. However, due to the reductionist nature of the extracted features, even measures incorporating many such features will potentially fail to capture biologically important song information. Additionally, as with direct human assessment of song similarity, these approaches often require significant human intervention for the selection of which samples of song material to analyze and which features to weigh in assessing similarity.

For these reasons, we were interested in developing an approach to scoring the similarity between pairs of songs (and other complex stimuli) that could reproduce some of the human capacity for holistically integrating across complex stimulus dimensions but that was also automatic, reproducible and efficiently deployed across large data sets. Our approach has four main components. First, we start with a representation of song that is the set of power spectral densities (PSDs) associated with each discrete syllable. This representation retains much of the complexity of the spectro-temporal information present in each song without specifically extracting features such as the pitch or entropy of each syllable. Second, we represent the spectro-temporal structure of each song by transforming each of a large number of individual syllables drawn from the song (as represented by their corresponding PSDs) into a ‘syllable similarity’ space, in which each syllable (PSD) is represented by its similarity to a large basis set of other syllables (PSDs). Intuitively, in this high dimensional space, different iterations of a given syllable type (e.g. different iterations of the syllable ‘A’) will cluster near each other, because they share a similar PSD. Each song is therefore associated with regions of high density within the syllable similarity space, with each high density region corresponding approximately to a distinct syllable type. Third, we then model each song by characterizing the high-density regions within the syllable similarity space (corresponding to regions in which there is clustering of PSDs). We use a Gaussian Mixture Model (GMM) to fit these regions of high density, in which the means and variances are fit to the data for each song using a standard expectation-maximization algorithm [14,15], and the number of Gaussian mixture components is determined using Bayesian Information Criteria (BIC)[16], a measure of model fit that is penalized for increasing model complexity. Lastly, since the GMM describing the distribution of spectrotemporal structure for syllables from each song is a statistical model, any differences in vocal structure (e.g. between tutors and tutees, or before and after onset of manipulations) can be assessed according to a principled information theoretic measure of the differences between the models fit to those songs. We describe the difference in structure between songs as the difference between two cross entropies, here referred to as the contrast entropy (CE, see Methods). CE quantifies the amount of information present in a reference song (e.g. a tutor song, or baseline song before a manipulation such as deafening) that is not present in a comparison song (e.g. the learned song of a tutee, or songs that are produced following deafening).

In this paper, we validate our approach by characterizing the similarities between pairs of songs for two conditions. First, we assess song learning by comparing adult tutor songs and juvenile tutee songs in the Bengalese finch (*Lonchura striata domestica*), a species with variability in both spectral content and syntax. We show that CE provides a measure of the quality of song learning in Bengalese finches that is well correlated with scores provided by human experts. Second, we assess song deterioration following deafening by comparing baseline songs produced by adult Zebra finches (*Taeniopygia guttata guttata*) with songs of the same individuals at varying times following deafening. We show that CE detects and quantifies the characteristic song deterioration that follows deafening[7]. Together, these results demonstrate an unsupervised approach to the quantification of both song learning and song deterioration. Such an approach allows holistic and reproducible, high-resolution tracking of song similarity across both large populations of individuals and long periods of time, and has potential relevance outside of birdsong as it could be applied to automatic analysis of human speech and other complex multidimensional data.

## Results

We present an automated method for assessing song learning by calculating the amount of spectral information which is present in the song of the tutor bird, but absent from the song of the tutee. To accomplish this we construct statistical models from the song spectral content of both reference song (tutor) and comparison song (tutee) then estimate the amount of reference song information accounted for by the tutor model but not the tutee model. Below, we first describe how the statistical model for each song is constructed and how the statistical models for two songs are compared to provide an estimate of song similarity, the contrast entropy (CE). We then show that CE correlates well with human scores for data sets that characterize song learning and demonstrate that CE also detects the typical deterioration of song following deafening. The descriptions below elaborate a specific instantiation of our overall approach. We follow this with an examination of the robustness of CE measures across a range of different possible instantiations, and a discussion of how our approach could be modified or extended to address related issues of similarity in other domains.

### Overview of statistical model assembly

The assembly of statistical models representing song spectral data is schematized in Figure 2. Starting with song data from a given bird (Fig. 2A), we identify individual syllables as continuous amplitude traces above a threshold. For each syllable, we calculate the power-spectral density (PSD; an estimate of acoustic power at a set of specific frequency values) using Welch’s method[17] (Fig. 2B). We then calculate the similarity between each PSD and a “basis set” of PSDs (see Methods), where the basis set is a random draw from the set of PSDs being modeled. These computed similarities create an NxM matrix of PSD similarities (Fig. 2C) where N is the number of PSDs being analyzed (’target’ PSDs) and M is the number of PSDs in the basis set (’basis’ PSDs). For the analyses presented below, we use a value for ‘N’ of 3000 PSDs drawn from the song to be modeled, and a value for ‘M’ of 50 PSDs to form the basis set for construction of the syllable similarity matrix.

**Fig 2.**
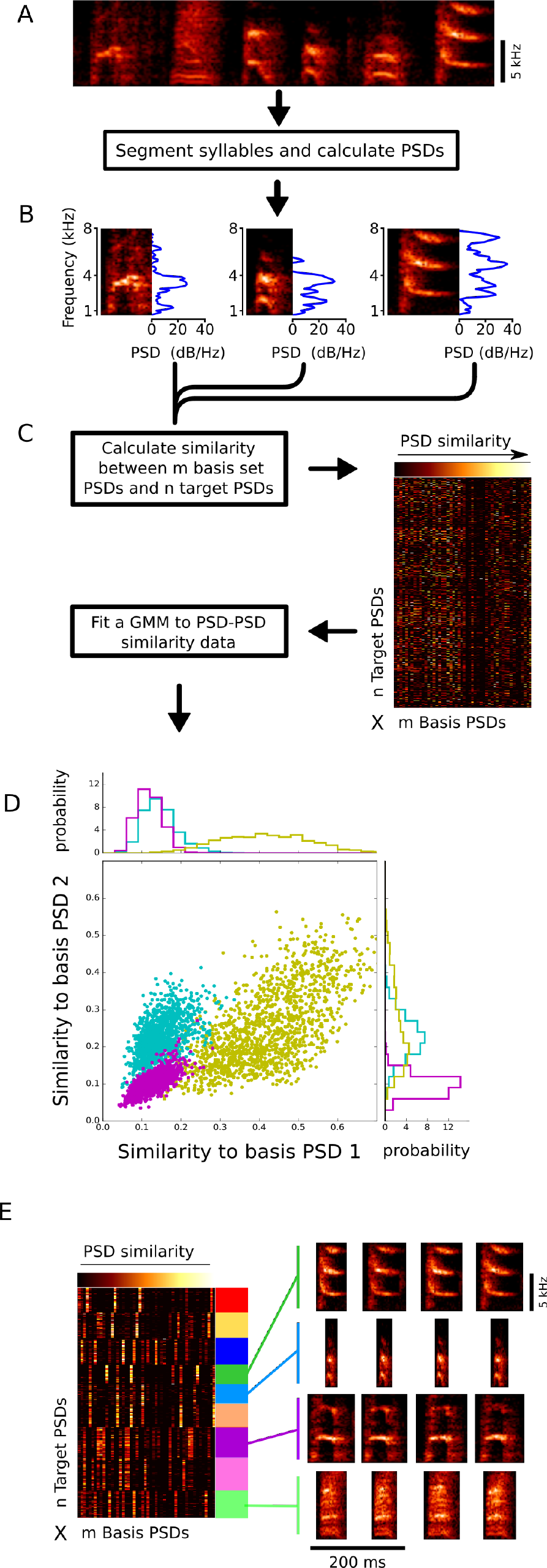
Assembly of statistical models for song. (A) To assemble a statistical model for the song of a given bird, we first segment all syllables from a set of songs produced by that bird and compute their corresponding PSDs. (B) Three examples of segmented syllables, each of a different type, and their corresponding PSDs. (C) For each of 3000 ‘target’ syllables from the song to be modeled, similarity of PSDs is calculated relative to a basis set of PSDs for 50 syllables randomly drawn from the same song. This creates an M (number of basis syllables) by N (number of target syllables) similarity matrix. (D) Visualization of how transformation of raw syllable data into the syllable similarity matrix results in a clustering of syllables by type. Each point in the plot indicates the similarity between the PSD for one target syllable and two basis PSDs (‘basis PSD1’ and ‘basis PSD2’) from the set of 50 basis PSDs. For clarity of exposition, only data that fall into one of three regions of high density are plotted here. Each of these regions corresponds approximately to multiple instances of one syllable type (which cluster near each other because of the similarity in their PSDs). In practice, there were more than three regions of syllable clustering (corresponding approximately to the number distinct syllable types in the bird’s song), and these regions were represented in the 50 dimensional space defined by the basis set of PSDs (only two of which are illustrated here). The regions of high density in this similarity space were fit with a Gaussian mixture model, in which the optimal number of Gaussian mixtures was determined by Bayesian Information Criteria. Individual data points here are color-coded by their assignment to one of three Gaussian mixtures, with data points corresponding to 6 additional Gaussian mixtures not shown. In any single dimension (top and right) data points assigned to each Gaussian mixture were approximately normally distributed. (E) Similarity matrix shown in C, reordered so that data are grouped by assignment to each of 9 Gaussian mixtures fit to the data (represented by colored blocks at the right of the similarity matrix). In this reordered representation, it is apparent that syllables assigned to each Gaussian mixture have a shared ‘bar code’ reflecting a shared pattern of PSD similarity values relative to the basis PSDs. The spectrograms at the right illustrate that syllables assigned to a given Gaussian mixture tend to be of the same type.

Translation of the raw syllable data into a matrix of syllable-similarities has the important consequence that it naturally and automatically clusters syllables with similar PSDs into regions of high density (Fig. 2D, E). Each of these regions corresponds approximately to groups of syllables that would be identified by a human observer as belonging to a specific type. That is because each instance of a syllables type has a similar PSD, and therefore each of these instances will have a similar pattern of distances from each of the elements of the basis set. This transformation therefore results in a representation of song in syllable similarity space in which thousands of exemplar syllables are clustered into a small number of high density regions, corresponding approximately to the numbers and identities of categorically distinct syllable types in the original song.

We then fit a series of GMMs[14] (using expectation maximization[15]) to the distribution of similarity values. Each successive model in the series has an incrementally higher number of mixture components. The model with the best fit to the data based on the lowest Bayesian Information Criterion[16] is then used to represent the song of that bird. Conceptually, the number of Gaussian mixture components in the song model corresponds approximately to the number of discrete syllable types present in the song.

The distribution of similarity scores reveals GMMs to be appropriate and effective for modeling these data. To illustrate this, Figure 2D depicts the similarities between three discrete target PSDs (Fig. 2D, blue purple and yellow dots) and two basis PSDs (out of a total of 50 basis PSDs). Target PSD identity was assigned as the maximum posterior probability over Gaussian mixture (see Methods) and is depicted by data color. Examination of the joint distribution of similarities between target PSDs and basis PSDs (Fig. 2D) reveals three clusters that are, even in just two dimensions, well separated. Importantly, the separation between clusters is reliant not only on the mean and variance in any one dimension but also on the covariance structure between values in the two exemplar dimensions; all three PSD categories are distributed elliptically. Examination of the marginal distributions (Fig. 2D, top and right) in each single dimension exemplifies the Gaussian nature of the data. Sorting the NxM matrix of similarity data by PSD identity (Fig. 2E) reveals the underlying ordered structure of these data; the PSDs in each group (Fig. 2E, indicated by color) share stereotypical PSD similarity structure. Spectrograms reveal good consistency of syllable identity (as indicated by corresponding PSD identity) within GMM classified groups (Fig. 2E, right).

### Assessment of the quality of learning

Because we capture song spectral content as statistical models, we can compare the spectral content of two songs using a principled information theoretic measure. In Figure 3 we outline the calculation of this measure, here called contrast entropy (CE). CE estimates the amount of information in the reference song that is not present in the comparison song (Fig. 3A). In this example, some, but not all, syllables from the reference song are well-represented in the comparison song. GMMs are fit independently to the two songs, as described above, except a single basis set of PSDs, drawn from the reference song, is used for modeling both songs, so that the syllables from both songs are transformed into the same syllable similarity space. In Figure 3B a single dimension of these probability densities are shown for simplicity. We then calculate the mean-log likelihood of a set of reference song data (Fig. 3C, light blue histogram). This specific data set was not used to estimate GMM parameter values (held-out data). This provides an estimate of the cross entropy between the sampled distribution of the reference song and the model for the reference song. We similarly estimate the cross entropy between the sampled reference song distribution and the model for the comparison song. Here we define the difference between these two cross entropies as the contrast entropy (CE).

**Fig 3.**
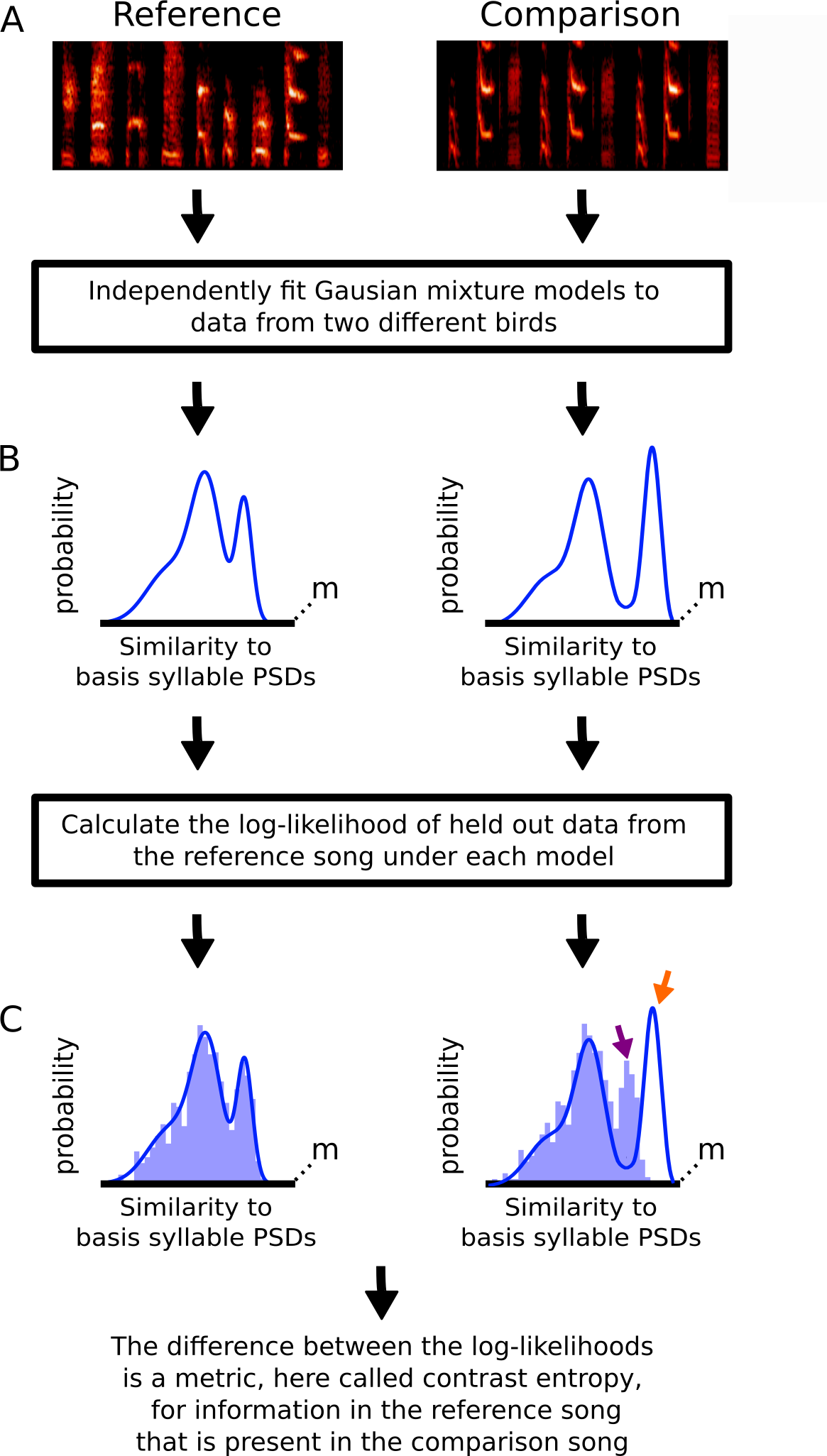
Estimation of the amount of spectral information present in the reference (tutor) song that is absent from the comparison (tutee) song. (A) Example reference and comparison songs. To compute contrast entropy for these songs, we first fit GMMs to the data from each song. (B) Representation in one dimension of the GMMs fit to song spectral content for both the reference song (left) and the comparison song (right). (C) We then calculate the log-likelihood of reference song syllables withheld from model fitting (light blue histogram) under both the model for the reference song and the model for the comparison song. Spectral content present in the reference song and not well represented in the comparison song model (purple arrow) will result in lower likelihood. Consequently, the difference (CE) between the two (reference song and comparison song) log-likelihoods will increase if there is information present in the reference song that is absent from the comparison song. However, information present in the comparison song, but not in the reference song (orange arrow) will not impact the CE.

An intuition for the relevance of CE to the content of song is provided by examination of Figure 3C. Here, represented in a single dimension, the held-out data from the reference song fit the probability distribution from the reference model well; very little of the data (histogram) fall outside the model (blue line). The same data are less well matched to the probability distribution for the comparison song; some data fall outside the model (Fig. 3C, purple arrow). Thus, the log-likelihood of these data given the reference song model is higher than that of the same data given the comparison song model and the magnitude of this difference determines the CE. However, the likelihood of the data is not influenced by portions of the distribution of the comparison song model that are not occupied (red arrow) by data from the reference song. Hence, contrast entropy will be low if the tutee (comparison) learned all elements of the tutor (reference) song well, but will be unaffected by tutee song content that is not present in the tutor song (e.g. song content innovated by the tutee; we discuss later how our approach can be extended to quantify innovation by the tutee).

### CE closely parallels human assessment of learning outcomes

We evaluated CE as a holistic measure of song similarity through comparison of CE to human scoring (Fig. 4). We used both CE and human scoring to assess learning in five cohorts of Bengalese finches. Each cohort was tutored with a different song (cohort tutor song). Figure 4A shows an example of the cohort tutor song for one group (top) and 5 tutee songs that illustrate a broad range in the quality of copying. For each cohort, four expert human evaluators independently estimated the similarity between each tutee’s learned song and the tutor song on a scale of 0-4, with 0 being most similar.

**Fig 4.**
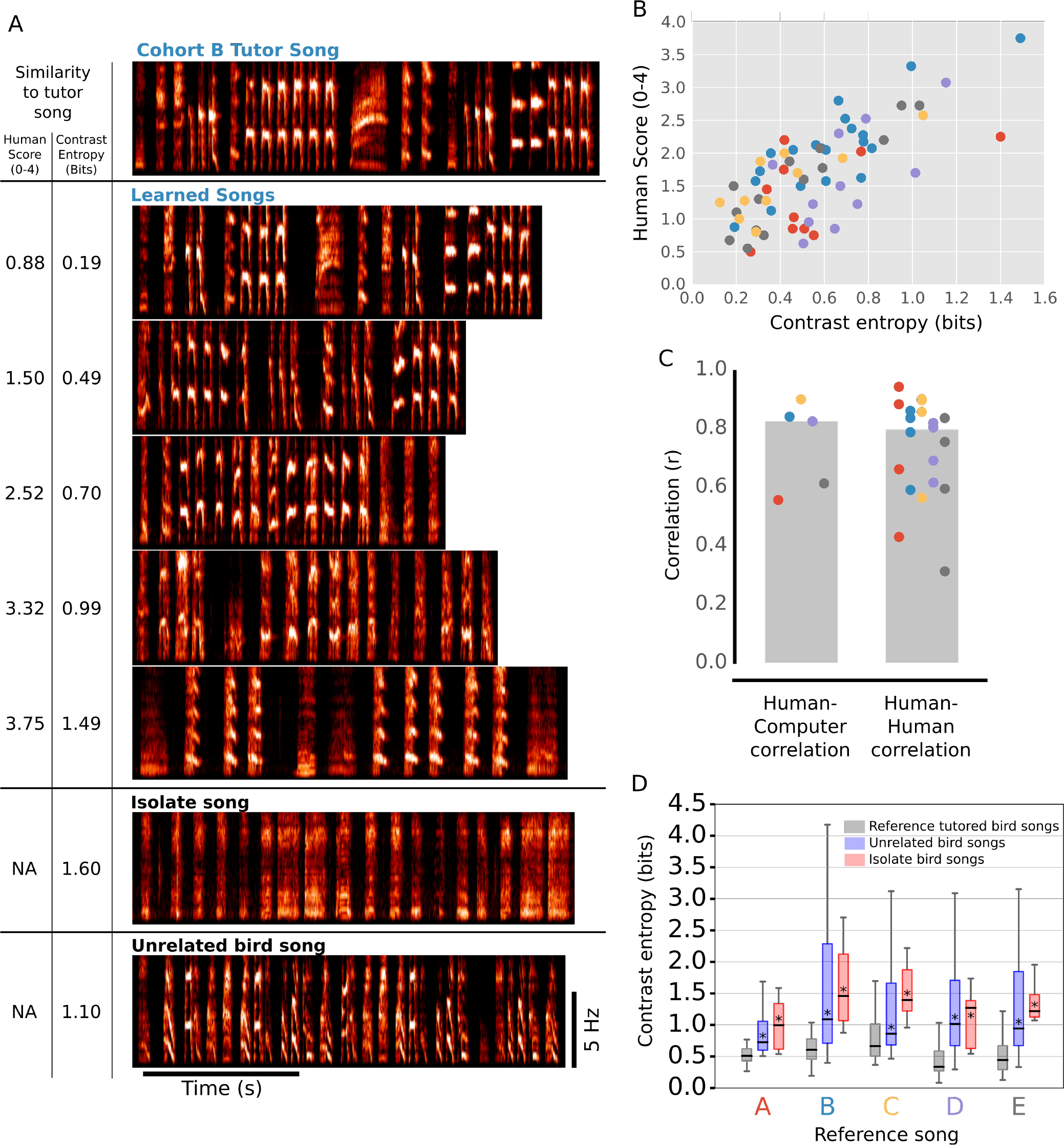
Contrast entropy closely parallels human assessment of learning outcomes. The quality of learning for individuals from five cohorts, each with a distinct tutor song, were evaluated by contrast entropy (CE) and human inspection. (A) Example spectrograms of the tutor song from one cohort and the songs of 5 tutees from the same cohort (cohort B). Also shown for comparison is the song of one isolate bird, raised without tutor song exposure (isolate song), and the song from one bird raised with a different tutor (unrelated bird song). Numbers at left indicate the CE and human similarity scores for each song relative to the tutor song from cohort B. (B) There was a good correspondence between CE and human evaluations of learning across a broad range. Here, human scores are the average of four human judges. Across all five cohorts, CE and human scores were well correlated (p < 0.01, r = 0.722, OLS). (C) Comparison of CE and human scores for each of the five cohorts. Human-computer correlation (left) shows the correlation between CE values and average human scores for each of the five cohorts. Human-human correlation (right) indicates the correlation between the scores of each of 4 individual humans and the average of the remaining human scores for each cohort. Medians are indicated as gray bars. (D) Summary of CE scores for the five cohorts (gray) were significantly lower than scores from a cohort of unrelated birds (purple, p < 0.01, Wilcoxon rank test) and from a cohort of ‘isolate birds’ raised without a tutor (red, p < 0.01, Wilcoxon rank test). Across all panels, bird cohort identity is indicated by color.

Across all five groups, CE and human scores were well correlated. Figure 4A shows the average human and CE scores assigned to 5 example tutee songs from one cohort. The rank ordering of similarities for these five songs relative to the tutor song was the same for CE and human scores; the learned songs are displayed from top to bottom in order of decreasing similarity to the tutor song by both measures. Figure 4B shows the correlation between CE and average human scores for 65 birds from the 5 cohorts (r=0.72, p<0.01). When calculated for each cohort individually, the median CE-human correlation was high (Fig. 4C,human-computor correlation). We compared these CE-human correlations to Human-Human correlations. For each cohort of birds, the scores of each evaluator were correlated with the average scores provided by the other evaluators (Fig. 4C, human-human correlation). The correlation between CE and human scores was comparable to the correlation between different humans’ scores. Together, these results indicate that CE provides a holistic and automated assessment of song learning that closely parallels human evaluation.

As an additional reference for ‘poor learning’, we also computed CE relative to each cohort tutor song for two additional groups of birds: ‘isolate birds’ that were raised without exposure to any tutor, and ‘unrelated birds’ that were raised with a tutor different from any of the five cohort tutors. CE indicates information from the reference song that is missing from comparison songs. Consistent with this, CE for isolate songs that contain atypical vocalizations was higher than that for songs from normally tutored birds (Fig. 4A, example ‘isolate song’ and Fig 4D, summary comparisons, p<0.01, Wilcoxon rank test). Similarly, CE for songs from birds that copied an unrelated tutor was also higher than that for songs from birds that learned from cohort tutor (Fig. 4A, example ‘unrelated song’ and Fig. 4D, summary comparisons; p<0.01, Wilcoxon rank test).

### Quantification of changes in the spectral content of song due to deafening

Many studies assess changes in song structure following various manipulations. One such manipulation, deafening, produces gradual deterioration of song structure and has been used extensively to provide insights into the mechanisms of song learning[6,7,18]. However, as with learning, quantitative assessment of song deterioration following deafening has often relied on human inspection[6,18]. To determine whether CE captures this deterioration we examined the songs of seven Zebra finches before and after deafening. We used CE to evaluate changes in song spectral content between baseline reference songs (before deafening) and comparison songs from the same birds two, four, six, and eight weeks post deafening. Figure 5A illustrates spectrograms from baseline songs and corresponding portions of post-deafening songs for 3 birds that qualitatively exhibited small (green), medium (yellow) and large (blue) changes to syllable spectral content. Figure 5B shows post deafening CE trajectories for 9 birds. These data reveal a gradual and continuous loss of song information over time (as quantified by CE), and demonstrate our approach as a sensitive method for evaluating changes in song following manipulations such as deafening.

**Fig 5.**
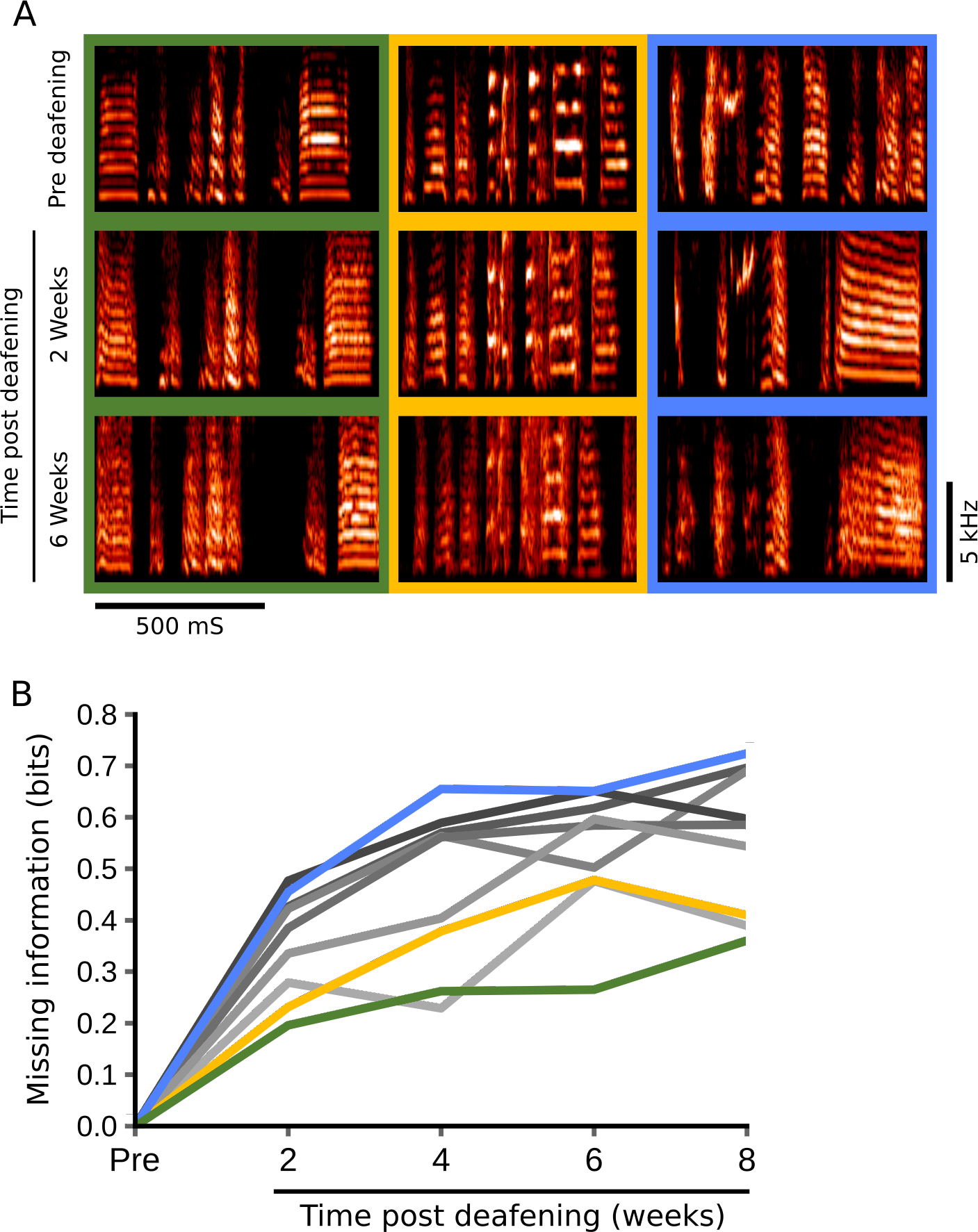
Quantification of changes to song following deafening. (A) Spectrograms from before (pre), two weeks, and six weeks post deafening for three Zebra finches demonstrate typical disruption to the spectral content of song due to deafening. (B) CE values indicating information missing from post deafening songs relative to baseline reference songs for nine birds at two, four, six, and eight weeks following deafening. Colors indicate bird identity, with green, yellow and blue in panels A and B illustrating data from birds that had small, intermediate and large changes to song spectral structure following deafening.

### Robustness of similarity measures to parameter choices

Our approach is intended to provide a measure of song similarity that requires little in the way of user intervention and selection of parameters. Correspondingly, all of the foregoing analyses were based on a specific instantiation of our approach using fixed values for parameters that could in principle be set by a user. Here we consider how different choices of parameter values affect similarity measures and demonstrate that CE measures are indeed very robust across a brand range of values. The specific parameters that we consider are 1) the number of ‘target’ syllables drawn from both the reference song and the comparison song, 2) the number of ‘basis’ syllables used for the PSD similarity basis set, and 3) the number of mixture components in the GMM used to model the structure of each song. In addition to these numerical choices, our approach as described above uses the PSD for each syllable as a representation of the spectro-temporal complexity of song. We therefore also consider and discuss below how different choices of song representation could affect similarity measures or extend our approach to capture other aspects of song structure.

There are two numerical values important to CE that are necessarily under experimenter control: the number of target syllables from the song to be modeled (N) and the number of syllables in the basis set (M). To determine appropriate values, we conducted numerical titrations of both the number of target syllables in the input data set (Fig. 6a and S1) and the number of syllables in the basis set (Fig. 6b and S2). For each of 44 birds, CE (relative to the corresponding tutor song) was calculated using 100, 200, 500, 1000, 2000, and 3000 target syllables from the songs that were being modeled (both reference and comparison songs). For each number of target syllables, we computed CE values for the 44 birds and correlated these CE values with CE values determined with 3000 syllables. Not surprisingly, CE values with little input data showed substantial deviation from the 3000 syllable CE values (Fig. 6a and S1A). However, for target syllable numbers above ~500, CE-CE correlations approached an asymptote, indicating little change to CE similarity measures above this value (Fig. 6a and S1B-D). We therefore used a fixed value of 3000 target syllables to model song structure throughout our analysis. For the same set of birds, CE values were calculated using 5, 10, 20, 40, 80, and 160 basis syllables. CE-CE correlations were calculated relative to CE values derived using the 160 PSD basis set. CE-CE correlations approached an asymptote for basis set numbers above ~25 (Fig. 6b and S2). For computational efficiency and to constrain the number of free parameters in our GMMs, we therefore used a basis set of 50 syllables throughout the study. These data indicate that CE measures of song similarity are robust to changes in numbers of target and basis syllables above threshold minimum values and therefore that out approach can be deployed effectively with fixed values of these parameters that do not require human tuning.

**Fig 6.**
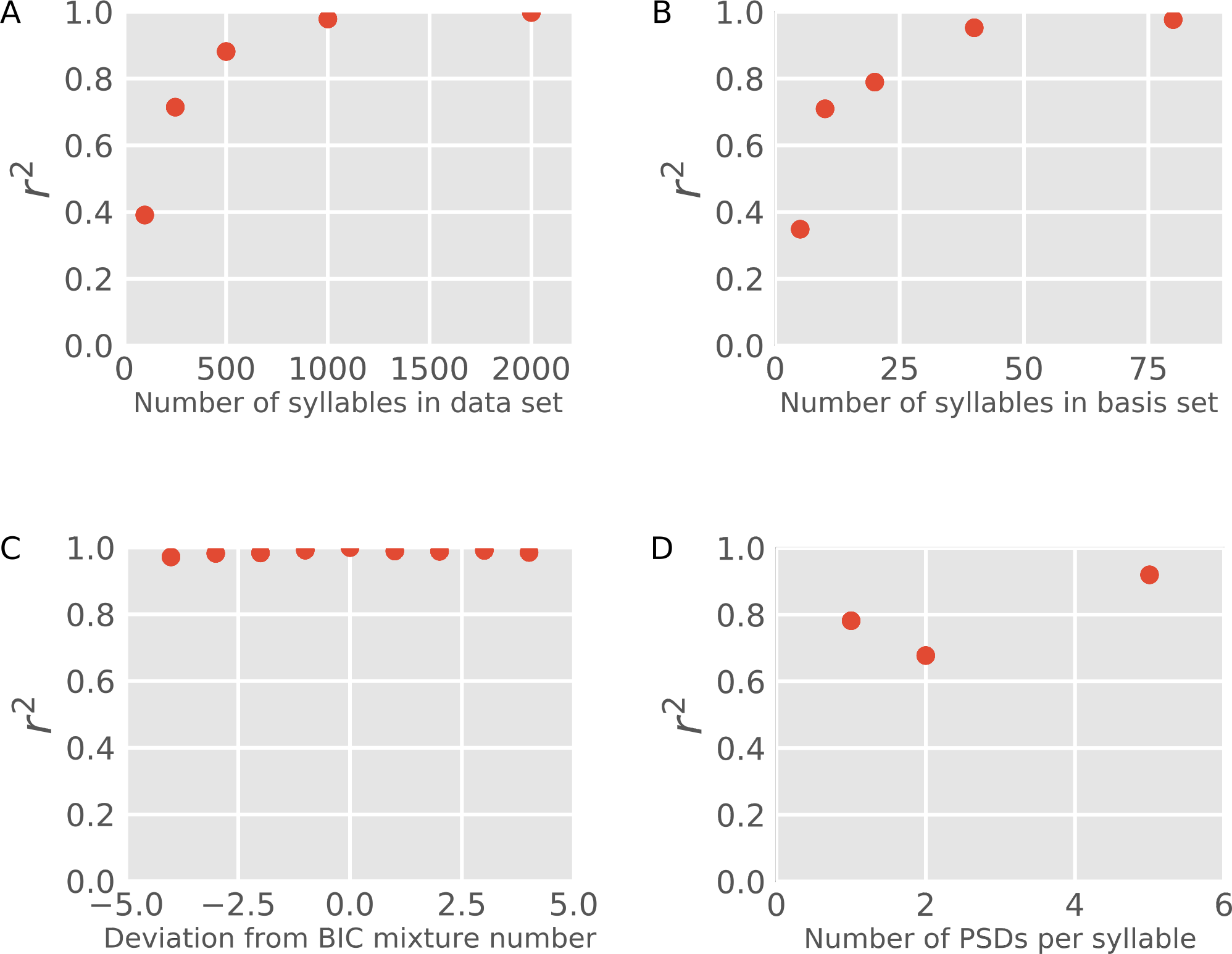
Contrast entropy song similarity measures are robust to variation in defined parameters. (A) Plot of r^2^ values for correlations between CE calculated using a range of input data sizes and CE calculated using 3000 syllables of input data. (B) Plot of r^2^ values for correlations between CE calculated using a range of basis set sizes and CE calculated using a 160 syllable basis set. (C) Plot of r^2^ values for correlations between CE calculated using the number of mixture components (k) determined by BIC (nBIC) and CE calculated using a number of mixture components ranging from nBIC-4 to nBIC+4. (D) Plot of r^2^ values for correlations between CE calculated using a single PSD representation of a syllable and CE calculated using a 10 PSD representation.

The number of Gaussian mixtures required to model the spectral complexity of a given song was selected automatically using Bayesian Information Criteria. However, the number of Gaussian mixtures can in principle be set to different values, for example in cases where the experimenter has an independent basis for modeling a song with a specific number of discrete syllable types. We therefore evaluated the robustness of CE similarity measures to variation in the number of Gaussian mixture components. Specifically, we calculated the squared error between CE values when calculated using the number of mixture components determined by BIC (nBIC) and CE values calculated using a series of other models in which the number of mixture components ranged from nBIC-4 through nBIC+4 (Fig. 6C and S3). These values were used in both the reference model and the comparison model. CE was very robust to variation in the number of mixture components; squared errors for all CE comparisons were above 0.96 (Fig. 6C, n=44, p<0.001 for all correlations).

Throughout the analyses described above, we used a single PSD to capture the spectro-temporal content of a given syllable. At each frequency represented, this single PSD encodes acoustic power averaged across the duration of the entire syllable (Fig. 2B) effectively capturing syllable spectral content collapsed across time. Accordingly, time dependent spectral information is not captured by these representations and this missing information may influence CE values. We therefore compared CE calculated using single PSDs per syllable with a series of CE values calculated using multiple PSDs per syllable where each syllable was divided into equal duration blocks (2, 5, or 10 blocks per syllable) and PSDs were calculated for each block (Fig. 6D and S4). CE calculated with single PSD syllable representations was well correlated with CE calculated with 10 PSDs per syllable (Fig. 6D and S4A; n=44, r^2^=0.78). This strong correlation suggests that much of the spectro-temporal information in a syllable is captured in single PSD representations. For purposes of computational efficiency, we therefore use a single PSD to represent syllable spectral content for song modeling and calculation of CE.

### Syllable identity assignments provided by GMMs are well correlated with human assignments

Our contrast entropy measure is intended primarily to provide a holistic measure of song similarity between a reference song and comparison song. As part of this process we model the structure of each song using a GMM in which an intuition is that each fit Gaussian corresponds approximately to what a human observer would label as a single categorically defined syllable type. The assignment of syllables to specific Gaussian mixtures is not required for the computation of CE, which relies solely on the models fit to the distribution of (unlabeled) syllable similarity data. However, it is of potential interest to know how effectively assignment of syllables to different Gaussian mixtures results in a categorization of syllables by type that matches human labeling of syllables. Such automatic labeling of syllables has potential utility in objectively determining the number of distinct syllable types in a bird’s song repertoire and facilitating the assignment of labels corresponding to these types to large amounts of song data.

Here, we explicitly examine the correspondence between syllable labels assigned using the GMM for each song to labels assigned by expert human evaluators. For each of 90 birds, syllables were labeled by determining the maximum posterior probability for assignment of syllable identity under the GMM fitted to songs of that bird (see Methods). These classifications were then compared with classifications provided by human inspection. Figure 7A illustrates labeled syllable categories for songs from two example birds. For some birds (e.g. upper panel) there was perfect concordance between human assigned (black) and GMM assigned (red) labels (categories). However, for most birds there were some discrepancies between human and GMM assigned labels (e.g. lower panel, gray box). To determine the accuracy of GMM based classifications, for 90 birds, all assignments were inspected by an expert human observer and the classification of each was determined to be accurate or inaccurate (Fig. 7B). Overall, 50% of birds had 96% or better correspondence between human and GMM assignments, while 80% of birds had 93% or better correspondence (Fig. 7C). This correlation between human and GMM based syllable classification indicates that much of the complex information subjectively used by humans to classify song syllables is incorporated into the GMM models that were used to model song structure.

**Fig 7.**
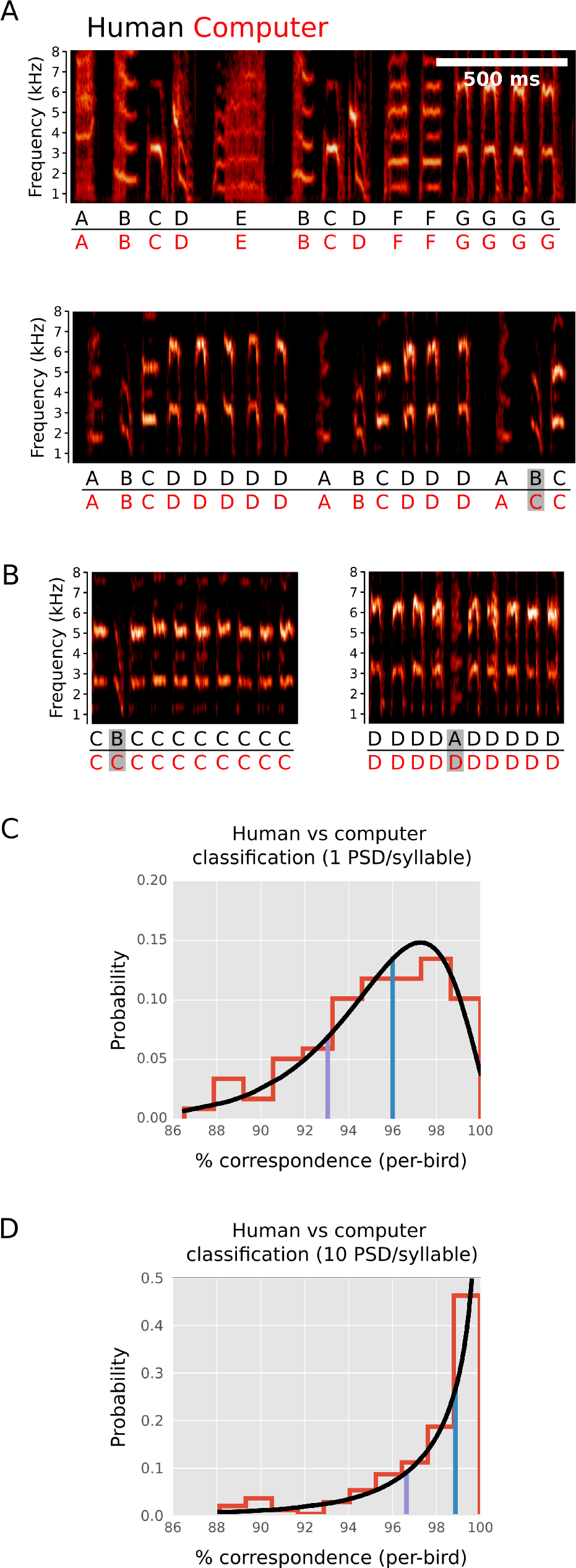
GMM derived syllable classifications are correlated with human syllable classifications. (A) Examples of labels assigned to two songs by human inspection (black) and GMM (red). For many birds, there were no differences between human assigned and GMM assigned labels (e.g. upper panel). However, for some birds, there were discrepancies (e.g. gray box, lower panel). (B) Erroneous GMM classifications can be identified by inspection of spectrograms for groups of syllables assigned to a given Gaussian mixture. Illustrated here are two examples of groups of syllables assigned to individual Gaussian mixtures, where it is apparent in each case that a single syllable (gray boxes) is miss classified relative to human assignment. For 90 animals, the number of miss-classified syllables was determined by such human inspection of groups of syllables that were assigned to each Gaussian mixture. (C) Distribution of the percent of correctly classified syllables (per-bird) is shown in red with a gamma distribution fit to these data shown in black. 50% of animals had greater than 96% correctly classified syllables (blue line) while 80% had more than 93% correctly classified syllables (purple line). (D) Distribution of the percent of correctly classified syllables per bird is shown as in C, but here with categorization carried out in which the input representation of each syllable to the GMM includes 10 PSDs evenly spaced over the duration of the syllable, rather than a single PSD for the entire syllable. Using this richer representation of syllable, 50% of animals had more than 99% correctly classified syllables (blue line) while 80% had more than 96% correctly classified syllables (purple line).

In this analysis, the spectral structure of each syllable was modeled using only a single PSD computed from the entire syllable (Fig. 2B). To ask whether a richer spectro-temporal representation of each syllable would increase concordance between human and GMM assignments, we built GMM models as above, but with 10 separate PSDs, evenly spaced over the duration of each syllable, used as an input representation for each syllable. We previously found that this richer syllable representation had little effect on CE measures of song similarity (Fig. 6D). In contrast, for the specific assignment of syllable labels, this richer representation resulted in significantly improved correspondence between human assignments and GMM assignments of syllable labels (Fig. 7D).

## Discussion

We demonstrate an approach for analysis of song and song learning that is computationally efficient and automated. We use syllable spectral content to assemble statistical models for song that can then be used to estimate the amount of spectral information present in one song but absent in another. Our measure of song similarity, the contrast entropy (CE), is well correlated with holistic song similarity scores provided by expert human evaluators while providing several critical advantages. CE is automatically computed and thus consistent given the same data, where human assessment is less reliable both across individuals and over time. Because CE is automatically and efficiently computed, we can analyze large amounts of data (many thousands of songs) facilitating dense analysis of learning across time and, for any given comparison, incorporating much more song data in assessments of learning than can be accomplished by a human evaluator. Importantly, this also obviates the need for selection of specific samples of song for comparison, and results in measurements of similarity that neither require, nor are potentially biased by, human intervention in selection of representative samples of song for analysis. Human evaluators of song learning also require species-specific training to ensure reliability and increase consistency across individuals. We show that our approach can be applied, with no modification, to two different songbird species vocalizations, indicating that this approach can be readily extend to analyze other vocalizations and other complex but similarly structured data.

CE is asymmetrical in that it estimates the amount of spectral content in a reference song that is missing from a comparison song. This asymmetry can be exploited to address distinct conceptual questions contingent on the reference-comparison relationship. In the case of birdsong, when the reference is tutor song, and the comparison is tutee song, the measure indicates how much information from the tutor song was not learned by the tutee. Reciprocally, if the reference is tutee song, and the comparison is tutor song, CE indicates how much information in the tutee song did not come from the tutor, providing an estimate of “innovation”.

We specifically focused on assessing the learning of song spectral content, but the general framework of comparing the shared information between statistical models can be extended (or restricted) to different categories of song information by changing the statistical descriptions of song. For example, an analysis might focus on the means of syllables but not the rendition-to-rendition variation. In this case, syllable variances would be excluded from cross entropy estimation. Alternatively, the model could be extended to include, not only spectral content, but also syllable transition information using a Hidden Markov Model (HMM) with Gaussian emissions. HMMs have been effectively used in the past to model song transition structure with human assigned syllable identities[19]. If song structure was modeled using HMMs with Gaussian emissions, CE would indicate discrepancies in spectral content as well as syllable ordering. Hence, our general approach to the evaluation of learning can be applied to any aspect of song that can be incorporated into a statistical model.

Beyond song learning, our approach allows fitting GMMs to sparse, high dimensional data. GMMs have been difficult to fit to high dimensional, sparse data, (like song PSDs) partially because standard marginalization based dimensionality reduction approaches (e.g. principle components analysis) remove covariance structure which is potentially critical to fitting accurate GMMs[20]. Here we reduce the dimensionality and the sparseness of our data via calculation of syllable-syllable similarities. Our results indicate that this intuitive approach allows modeling high dimensional data sets within a framework that facilitates quantitative and meaningful comparisons. Similarity matrices are already used in the context of spectral[21] and hierarchical[22] clustering, but neither approach provides a statistical description (provided by GMMs) of data and, thus, cannot be easily used for information theoretic calculations. Our approach may have broad application to high dimensional problems where statistical descriptions can be leveraged for more accurate classification or where information theoretic calculations are desired.

## Methods

### Song recordings

For audio recording, animals were single housed in sound isolation chambers (Acoustic Systems). Songs were recorded digitally at a sampling frequency of 32 kHz and a bit depth of 16 then stored uncompressed using custom Python or LabView (National Instruments) software. Recording microphones were placed in a fixed position at the top of the cage housing the bird. Prior to further analysis, all songs were digitally high pass filtered at 500 Hz using a digitally implemented elliptical infinite impulse response filter with a passband edge frequency of 0.04 radians.

### Syllable segmentation

Discrete units of sound separated by silence (syllables) were identified based on amplitude. First an “amplitude envelope” was created by rectifying the song waveform then smoothing the waveform through convolution with an 8 ms square wave. This amplitude trace was then used, through thresholding, to identify periods of vocalization. To automatically identify a threshold capable of separating vocalizations from silence, we used Otsu’s method[23]. Briefly, Otsu’s method is an exhaustive search to identify a threshold that minimizes the shared variance between data above threshold and data below threshold. Once the threshold is established, “objects” are identified as contiguous regions of the amplitude envelope over threshold. To eliminate short and spurious threshold crossing that can occur at the edge of syllables where syllable amplitude is low, any objects separated by a gap of 5 ms or less are merged, and then any objects shorter than 10 ms are eliminated. The onsets and offsets of each object are padded by 3 ms and then used to segment audio data from the original filtered waveform. We refer to each of these segments of audio data as syllables.

### Power spectral density estimation

To estimate frequency information of syllables while removing temporal information, we calculated the power spectral density for each syllable at 2048 frequencies using Welch’s method[17]. Briefly, the PSD was computed via FFT for successive 4096 sample windows (at 32kHz), each overlapping by 256 samples. These were then averaged over the duration of the syllable, and the power in the frequency range 600 Hz – 1600 Hz (sampled at 1970 points) was used as the PSD for the syllable.

### Similarity matrix

Instead of clustering syllables on their PSD values (sampled at 1970 points), we transformed each PSD into a syllable similarity representation. For each of M syllables, the Euclidean squared distance between the PSD of that syllable and the PSDs of a basis set of N reference syllables was calculated creating an MxN distance matrix, *D*, in which *D_ij_* = ∥*p_i_ – q_j_*∥^2^ where *p* is the vector of M syllable PSDs and *q* is the vector of basis set syllable PSDs. We then calculated A, a similarity matrix, where *A_ij_*=1 / (*D_ij_ /* max(*D))*. For each bird in our main analysis presented in results, an M=3000 data syllables and N=50 reference basis syllables were used.

### Gaussian mixture model and parameter estimation

We model each song as a Gaussian mixture model fit to the distribution of syllables in the syllable similarity space. These GMMs[14] are defined as:

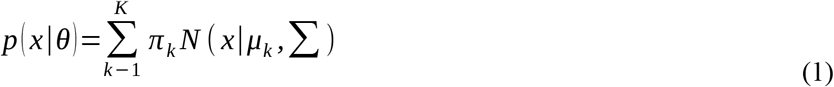

Where K is the number of mixing components, ***π_k_*** is the mixing weight, ***µ**_k_* is the vector of means, and ∑_*k*_ is the full covariance matrix for component *k*. The values of ***Ɵ*** ={ **π**_*k*_, **µ**_*k*_, **∑**_*k*_} are the parameters. The values of ***µ**_k_* and **∑**_*k*_ are initialized using the K-means algorithm and then estimated through standard expectation-maximization[15]. The value of π_*k*_ is *1/k* for all k. The values of *x* are the observed data. For this work, the implementation of expectation-maximization for GMMs in the scikit-learn software package[24] was used.

### Model estimation

We conducted model selection to identify the number of Gaussian mixtures needed to describe a song. For each bird, we fit, as described, a series of GMMs to song data. The set of models had increasing numbers of Gaussian components (K), ranging from 2-20. For each model we calculated a three-fold cross validated Bayesian information criterion (BIC)[16], a measure of model fit that is penalized for increasing model complexity. As the number of Gaussian components increase, the BIC decreases to a minimum value and then, as the number of Gaussian components proceeds beyond optimal, the BIC increases again. The number of Gaussian components in the model with the lowest BIC value was used to model each song’s structure for purposes of song similarity comparisons.

### Information theoretic measurement of song similarity

In order to quantify the similarity of two songs, we follow the procedure described above to derive two song models, one from a reference song (i.e. the song of an adult tutor, or baseline song of a bird prior to a manipulation such as deafening) and one from a comparison song (i.e. the song of a tutee, or song of a bird following a manipulation such as deafening). We then calculate how much worse (in bits of information) the model based on the comparison song is than the model based on the reference song at explaining held out data from the reference song. Mathematically, this is a difference between two cross-entropies (or Kullback-Leibler divergence). Here we refer to this quantity as contrast entropy, defined as follows:

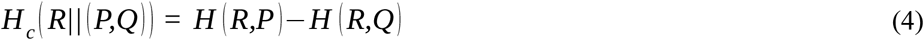

Where *H_c_* is the contrast entropy, *P* is the density function for the model fit to the reference data set, *Q* is the density function fit the comparison data set, *R* is the true (but unknown) density function of the reference data. *H(R,P)* is the cross entropy between *P* and *R*, and *H(R,Q)* is the cross entropy between *Q* and *R*. *H_c_* is calculated as:

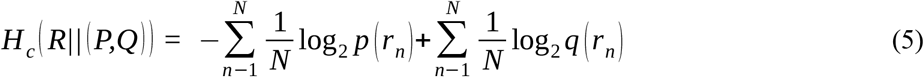

where N is the size of a set of syllable similarity data (*r*) from the reference song, ^log_2_*q*(*r_n_*)^ is the log likelihood of that data under the GMM (*p*) fit to a different set of reference data, and ^log_2_ *q* (*r_n_*)^ is the log likelihood of that data under the GMM (*p*) fit to data from the comparison song. Intuitively, *H_c_* is an estimate of how much worse the comparison (e.g. tutee) model is at explaining the true data from the reference song (e.g. tutor) than a model based on different (held out) data from the reference song. Alternatively, *H_c_* can be viewed as an estimate of the information present in the reference song that is absent in the comparison song.

### Syllable classification

For each syllable, the GMM based syllable classification was calculated as the maximum posterior probability over the K syllable types defined by each of the Gaussian mixtures in the fit model. Calculated as:

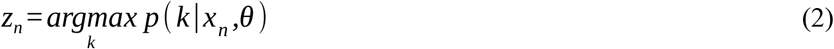

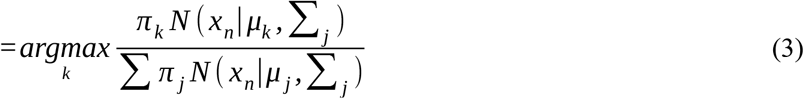

Where *z_n_* is the assigned syllable type.

### Human scoring

Contrast Entropy estimates of song similarity created by our method were compared with assessments made by humans experienced in song analysis. We considered 5 groups of songs, corresponding to 5 nests in our colony. Each group consisted of a tutor reference song (the adult male breeder in the nest) and multiple tutee comparison songs (between 8 and 20 tutee songs per group, corresponding to juveniles that were hatched and raised to independence in the nest of the adult male tutor). These groups were chosen such that each of the tutor songs were qualitatively distinct, and each group of tutees expressed a broad range in the quality of tutor song copying. Human judges that had extensive experience in song analysis were presented with spectrograms (frequency range from 500-10000 Hz) representing four samples of tutor song and four sample of each tutee song. Four seconds of each of the four sample songs were presented on the same time scale in a single representation. Human judges assigned a score between zero (high similarity to tutor), and four (low similarity to tutor), to each tutor-tutee song pair.

### Deafened birds

Deafening data were presented previously in Kojima et. al. 2013[24]. For each of seven birds, song was recorded before deafening and at two, four, six, and eight weeks post deafening. In each case the amount of information loss at any time post deafening was taken with reference to the pre-deafening, baseline song.

## Bibliography

1. Doupe AJ, Kuhl PK. Birdsong and human speech: common themes and mechanisms. Annu Rev Neurosci. 1999;22: 567–631. doi:10.1146/annurev.neuro.22.1.567

2. Brainard MS, Doupe AJ. Translating birdsong: songbirds as a model for basic and applied medical research. Annu Rev Neurosci. 2013;36: 489–517. doi:10.1146/annurev-neuro-060909-152826

3. Catchpole C, Slater PJB. Bird song: biological themes and variations. Cambridge University Press; 2003.

4. Tchernichovski O, Lints T, Mitra PP, Nottebohm F. Vocal imitation in zebra finches is inversely related to model abundance. Proc Natl Acad Sci U S A. 1999;96: 12901–4. Available: http://www.ncbi.nlm.nih.gov/pubmed/10536020

5. Thorpe WH. The Process of Song-Learning in the Chaffinch as Studied by Means of the Sound Spectrograph. Nature. 1954;173: 465–469. doi:10.1038/173465a0

6. Brainard MS, Doupe AJ. Interruption of a basal ganglia|[ndash]|forebrain circuit prevents plasticityof learned vocalizations. Nature. Nature Publishing Group; 2000;404: 762–766. doi:10.1038/35008083

7. Konishi M. The Role of Auditory Feedback in the Control of Vocalization in the White-Crowned Sparrow1. Z Tierpsychol. Blackwell Publishing Ltd; 1965;22: 770–783. doi:10.1111/J.1439-0310.1965.TB01688.X

8. Scharff C, Nottebohm F. A comparative study of the behavioral deficits following lesions of various parts of the zebra finch song system: implications for vocal learning. J Neurosci. 1991;11.

9. Tchernichovski, Nottebohm, Ho, Pesaran, Mitra. A procedure for an automated measurement of song similarity. Anim Behav. 2000;59: 1167–1176. doi:10.1006/anbe.1999.1416

10. Burkett ZD, Day NF, Peñagarikano O, Geschwind DH, White SA. VoICE: A semi-automated pipeline for standardizing vocal analysis across models. Sci Rep. Nature Publishing Group; 2015;5: 10237. doi:10.1038/srep10237

11. Wu W, Thompson J a, Bertram R, Johnson F. A statistical method for quantifying songbird phonology and syntax. J Neurosci Methods. 2008;174: 147–54. doi:10.1016/j.jneumeth.2008.06.033

12. Mandelblat-Cerf Y, Fee MS, Nottebohm F, Williams H, Marler P, Tchernichovski O, et al. An Automated Procedure for Evaluating Song Imitation. Bolhuis JJ, editor. PLoS One. Public Library of Science; 2014;9: e96484. doi:10.1371/journal.pone.0096484

13. Ho CE, Pesaran B, Fee MS, Mitra PP. Characterization of the structure and variability of zebra finch song elements. Proceedings of the joint symposium on neural computation. 1998. pp. 76–83.

14. Pearson K, Pearson A. Contributions to the Mathematical Theory of Evolution. Source Philos Trans R Soc London A. 1894;185: 71–110. Available: http://www.jstor.org/stable/90667

15. Dempster AP, Laird NM, Rubin DB. Maximum Likelihood from Incomplete Data via the EM Algorithm. Source J R Stat Soc Ser B J R Stat Soc Ser B. Methodological; 1977;39: 1–38. Available: http://www.jstor.org/stable/2984875

16. Schwarz G. Estimating the Dimension of a Model. Ann Stat. Institute of Mathematical Statistics; 1978;6: 461–464. doi:10.1214/aos/1176344136

17. Welch P. The use of fast Fourier transform for the estimation of power spectra: A method based on time averaging over short, modified periodograms. IEEE Trans Audio Electroacoust. 1967;15: 70–73. doi:10.1109/TAU.1967.1161901

18. Nordeen KW, Nordeen EJ. Auditory feedback is necessary for the maintenance of stereotyped song in adult zebra finches. Behav Neural Biol. 1992;57: 58–66. Available: http://www.ncbi.nlm.nih.gov/pubmed/1567334

19. Kogan JA, Margoliash D. Automated recognition of bird song elements from continuous recordings using dynamic time warping and hidden Markov models: A comparative study. http://dx.doi.org/101121/1421364. Acoustical Society of America; 1998; doi:10.1121/1.421364

20. Azizyan M, Singh A, Wasserman L. Efficient Sparse Clustering of High-Dimensional Non-spherical Gaussian Mixtures. 2014; Available: http://arxiv.org/abs/1406.2206

21. von Luxburg U. A Tutorial on Spectral Clustering. 2007; Available: http://arxiv.org/abs/0711.0189

22. Johnson SC. Hierarchical clustering schemes. Psychometrika. Springer-Verlag; 1967;32: 241–254. doi:10.1007/BF02289588

23. Otsu N. A Threshold Selection Method from Gray-Level Histograms. 1979;9: 62–66.

24. Pedregosa F, Varoquaux G, Gramfort A, Michel V, Thirion B, Grisel O, et al. Scikit-learn: Machine Learning in Python. J Mach Learn Res. 2011;12: 2825–2830.

